# Comparing current noise in biological and solid-state nanopores

**DOI:** 10.1101/866384

**Authors:** A. Fragasso, S. Schmid, C. Dekker

## Abstract

Nanopores bear great potential as single-molecule tools for bioanalytical sensing and sequencing, due to their exceptional sensing capabilities, high-throughput, and low cost. The detection principle relies on detecting small differences in the ionic current as biomolecules traverse the nanopore. A major bottleneck for the further progress of this technology is the noise that is present in the ionic current recordings, because it limits the signal-to-noise ratio and thereby the effective time resolution of the experiment. Here, we review the main types of noise at low and high frequencies and discuss the underlying physics. Moreover, we compare biological and solid-state nanopores in terms of the signal-to-noise ratio (SNR), the important figure of merit, by measuring free translocations of a short ssDNA through a selected set of nanopores under typical experimental conditions. We find that SiN_x_ solid-state nanopores provide the highest SNR, due to the large currents at which they can be operated and the relatively low noise at high frequencies. However, the real game-changer for many applications is a controlled slowdown of the translocation speed, which for MspA was shown to increase the SNR >160-fold. Finally, we discuss practical approaches for lowering the noise for optimal experimental performance and further development of the nanopore technology.

## Introduction

Nanopores are promising tools for biosensing applications and sequencing of DNA and proteins, as they can resolve single analyte molecules, resolve structural modifications of molecules, and even discriminate between nucleotide sequences^1–10^. The detection mechanism is simple: while passing through the pore, a (part of a) molecule transiently blocks the ionic current, thereby inducing a small dip in the current signal, which is detectable by the electronics (Fig. 1). The electrical read-out is carried out by an amplifier, which senses and amplifies the current signal, followed by a digitizer that performs the analog-to-digital conversion (ADC) of the data. Digital low-pass (LP) filtering is typically used to reduce the high-frequency noise, and thus improve the signal-to-noise ratio (SNR). Such a gain in SNR comes, however, at the expense of a lower time resolution, thereby imposing an inherent trade-off.

**Figure 1:**
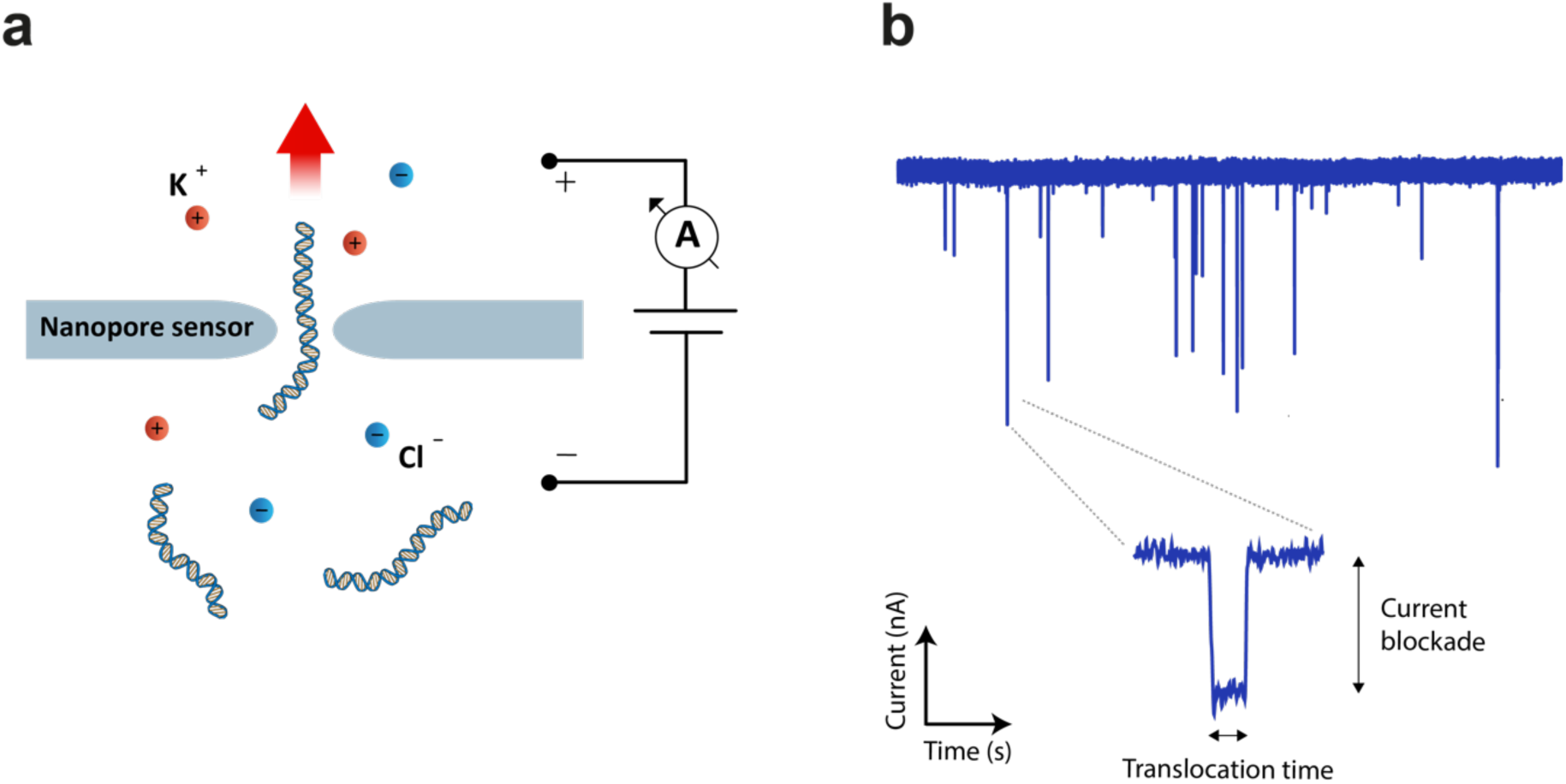
Fundamental principle of nanopore sensing. (a) A nanopore separates two aqueous compartments filled with electrolyte solution (e.g. potassium chloride) and small molecules (e.g. DNA) are electrostatically pulled through the pore by an applied potential. (b) While passing through the nanopore, the molecule temporarily induces a partial current blockade which is detected by an amplifier. The signature of a single-molecule translocation event is generally characterized by the amplitude of the current blockade, which is proportional to the volume of the molecule in the nanopore, and the translocation time, which represents the time that the molecule spends inside the pore.

The detection of analytes with nanopores thus is, on the one hand, limited by the ionic current noise which requires LP filtering that sets a finite operating bandwidth^11,12^, but on the other hand, by the fast speed (typically sub-milliseconds) at which molecules translocate through the pore, which conversely requires a high time resolution for accurate sampling. Various approaches have been investigated in order to slow down the molecular translocation. For biological nanopores, a DNA-translocating motor protein (such as a helicase or polymerase) has been used to slowly feed a ssDNA strand into a protein pore for DNA sequencing^13–15^. For solid-state nanopores fabricated in thin SiN_x_ membranes^16–18^ or 2D materials (graphene^19–21^, boron nitride^22–24^, molybdenum disulfide^25–27^), various efforts have been made to either increase time resolution^16,17,28–31^, or slow down the translocation process^32^ by the use of ionic liquids^27^, pore surface engineering^33^, mechanical manipulation with a double pore system^34^, and optical trapping^35^. Nevertheless, the SNR has not yet allowed *de novo* DNA sequencing with solid-state pores. An understanding of the noise sources that affect nanopore systems and how these govern the SNR is key for achieving signals wherein molecular structures can be resolved fast and reliably. Noise characteristics of nanopores have been reported in various isolated reports, but a systematic overview and comparison between biological and solid-state nanopores is lacking.

In this review, we first describe the typical noise sources that affect the ionic current recordings of biological and solid-state nanopores, both at low and high frequencies. Next, we compare their respective performances of various nanopores using ssDNA poly(dT) translocations as a test system. We assess the SNR under typical experimental conditions for different protein pores *Mycobacterium smegmatis* porin A (the M2 mutant with a neutral constriction and positively charged vestibule, subsequently referred to as MspA)^36^, *Staphylococcus aureus* alpha-hemolysin (α-HL)^37,38^, *Fragaceatoxin C* (the mutant of FraC with a positively charged constriction, referred to as ReFraC)^39,40^, and SiN_x_^29^ and MoS_2_^41^ solid state nanopores. We find that biological pores generally exhibit lower noise (Fig. 2a). Nevertheless, solid-state nanopores achieve the best SNR, largely because of the higher voltages and bandwidths that such devices can operate at, as compared to biological nanopores. Finally, we discuss approaches for lowering the ionic current noise and improving the SNR in biological and solid-state nanopores.

**Figure 2:**
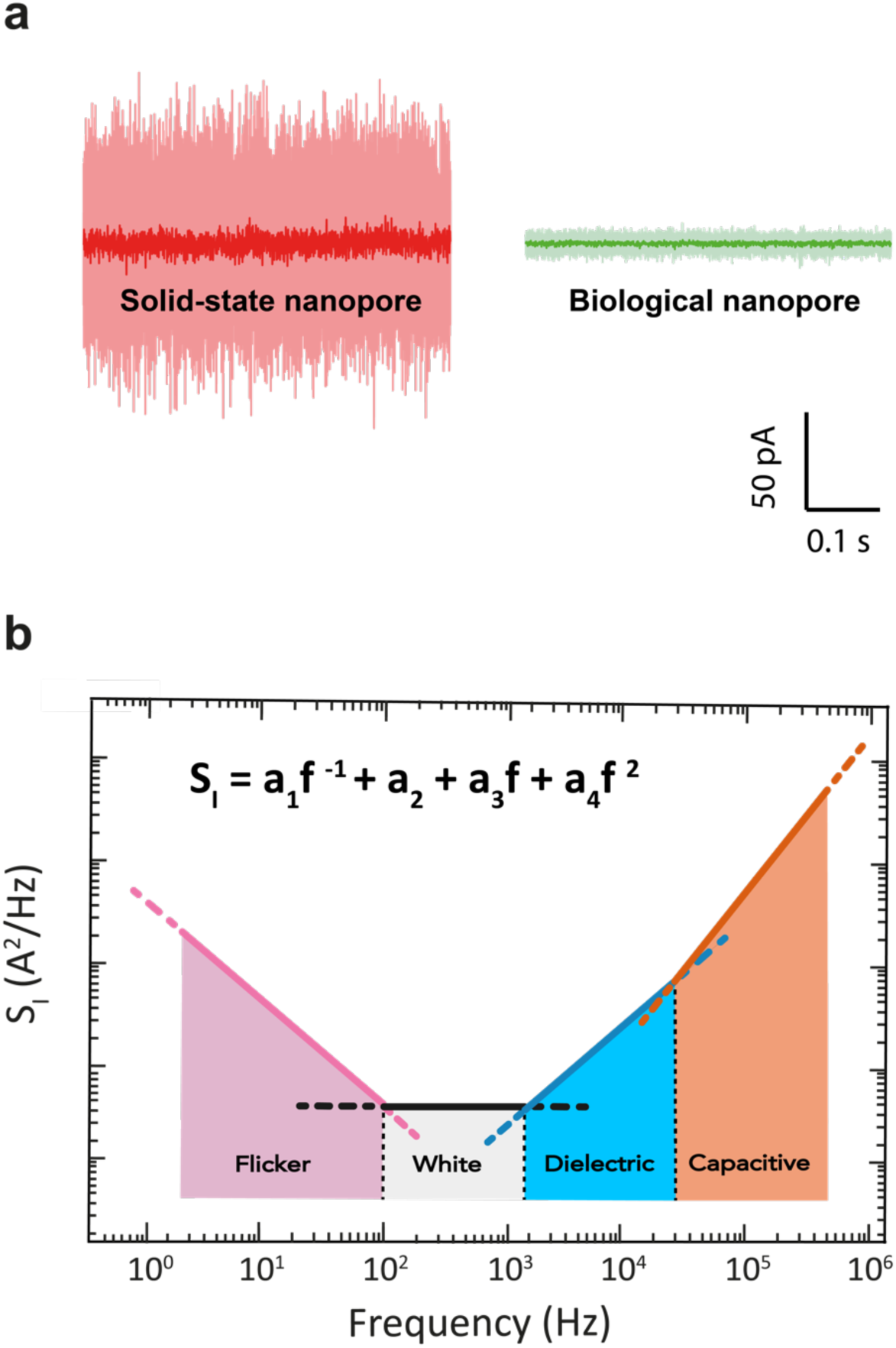
Ionic current noise in nanopores. (a) Example current traces for a 1.3 nm diameter solid-state SiN_x_ nanopore (red) and a 1.4 nm diameter biological *α*-HL pore (green), performed at a constant applied bias of 100 mV in 1 M KCl buffer at pH 7 at a bandwidth of 10 kHz (light) and 1 kHz (dark). (b) Schematic of the current Power Spectral Density (PSD) for a typical nanopore. Common types of noise are highlighted in the various frequency ranges.

### Noise sources in nanopores

Noise refers to any statistical fluctuation of a signal. It can be characterized by the standard deviation σ or root-mean-square (rms) variation around the average value as measured over the full bandwidth B of the signal, and by its power spectral density (PSD). Generally, noise is undesirable, as it can distort or even completely mask the actual signal. Nanopores typically operate by measuring a through-pore ionic current that is driven by a constant applied bias voltage. For the open-pore current measurement, where no analyte molecules are present, any deviation from the baseline current can be regarded as noise (Fig. 2a).

Understanding the origins of noise is fundamental for optimizing signal detection. Nanopore systems exhibit a range of different noise sources^42,43^. In Fig. 2b, we illustrate the major current noise sources that affect nanopore systems at different frequencies. Generally, these can be divided in: (i) low-frequency (≲100 Hz) 1/f noise and protonation noise; (ii) shot noise and thermal current noise (~0.1-2 kHz), which are both white noise sources (i.e., frequency-independent); (iii) high-frequency dielectric (~1-10 kHz) and (iv) capacitive (> 10 kHz) noise.

In the low-frequency range, 1/f noise (also referred to as ‘flicker’ or ‘pink’ noise) typically is the dominant source of noise. Its power decreases with frequency f following a 1/*f*^*β*^ scaling, with *β* ≈ 1. While this type of noise is found in many biological and physical systems, a fundamental understanding of it is still missing^44^. Based on phenomenological evidence, 1/f noise in nanopores has been associated with physical processes such as slow fluctuations in the number and mobility of the charge carriers^45–48^, nanometer-sized bubbles in the pore channel^49^, noise arising from the electrodes^50^, mechanical fluctuations of the freestanding membrane (e.g. for 2D materials)^23,51,52^, and conformational changes in the case of biological nanopores^53,54^. Smeets *et al* (2008)^55^ found that Hooge’s phenomenological formula^48^ could effectively describe the 1/f noise in solid-state^50,55–58^ nanopores,

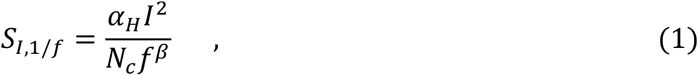

where Hooge’s constant *α*_*H*_ is an empirical parameter that quantifies the magnitude of 1/f noise fluctuations, *I* the ionic current, and *N*_*c*_ the number of charge carriers in the pore volume, which was further validated by follow-up studies^50,56–58^. As discussed below, solid-state nanopores typically feature a relatively pronounced 1/f noise, whose microscopic origin often remains unresolved. For biological pores, the low-frequency noise is typically dominated by protonation noise, which is generated by protonation/deprotonation of ionizable sites within the protein channel^59–61^. It can be described by fluctuations between two different current levels with mean lifetimes *τ*_1_ and *τ*_2_ for the protonated and deprotonated states, respectively, yielding a Lorentzian-shaped component in the frequency spectrum (for a complete derivation see Machlup et al., 1954^62^),

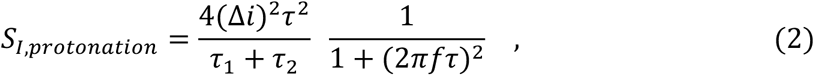

where ∆*i* is the difference in current between the two levels, and *τ* is the characteristic relaxation time, that can be expressed as *τ* = *τ*_1_*τ*_2_/(*τ*_1_ + *τ*_2_). For alpha-hemolysin, for example^60^, *τ* was found to be 3.1 × 10^−5^ *s*. A distribution of multiple Lorentzian processes such as in Eq. (2) can lead to 1/f noise^45^. Temporal conformational changes of the pore channel can also generate conductance fluctuations resulting in 1/f noise. Such a phenomenon, also known as ‘channel breathing’, was reported to affect protein pores such as bacterial porin channels^53,54^.

In the mid-frequency range (typically ~0.1-2 kHz), a frequency-independent white noise is observed that derives from thermal noise (also known as Johnson-Nyquist noise) and shot noise. Thermal current noise is fundamental to any dissipative element^63,64^ and adds to the current noise as

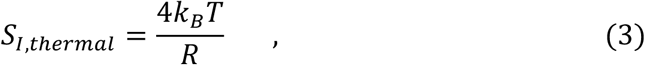

where *k*_*B*_ is the Boltzmann constant, *T* is temperature, and *R* the equivalent resistance of the nanopore. Shot noise, on the other hand, is due to the quantization of charge and is generated when charge carriers flow across a potential barrier^65,66^. Its current-dependent contribution to the noise can be expressed as

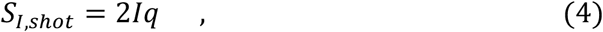

where *q* is the charge of a single carrier. In practice, shot noise and thermal noise are comparable in magnitude for the conditions that are typically used in nanopore experiments.

Another contribution to the nanopore noise is the thermal voltage noise generated by the loss conductance of the membrane and chip support that leads to dielectric noise^42,43^. As this conductance scales linearly with frequency, this noise can be described by

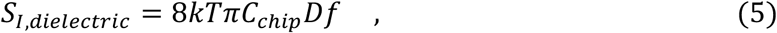

where *C*_*chip*_ is the parasitic capacitance and *D* a dissipation factor of the dielectric materials constituting the membrane and support chip. This source of noise typically dominates in the 2-10 kHz frequency range. To estimate *C*_*chip*_, one can simply use the expression for a parallel plate capacitor *C* = *εA*/*d*, where *ε* is the dielectric constant of the membrane material and *A* and *d* are the area and the thickness of the membrane, respectively. For *f* > 10 *kHz*, the current noise is determined by the input-referred thermal voltage noise *ν*_*n*_ across the total capacitance *C*_*tot*_ at the amplifier input^42,43^,

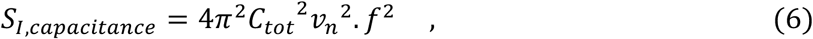

where *ν*_*n*_ is the input voltage noise (3 nV/Hz^−1^ for the commonly used amplifier Axopatch 200B^67^, Molecular Devices, San Jose, USA). *C*_*tot*_ is the total capacitance including the membrane and support chip capacitance *C*_*chip*_, the capacitance *C*_*amp*_ at the input of the amplifier, and the capacitance *C*_w_ of the wiring between the electronics and the pore. Notably, *S*_*I,capacitance*_ has an even stronger, *f*^2^, frequency dependence than *S*_*I,dielectric*_. The total current noise of a nanopore system over its full bandwidth is the sum of all contributions (Fig. 2b), i.e., the sum of Eqs. (1)–(6).

### Noise in biological nanopores

Biological nanopores are formed by the spontaneous insertion of membrane proteins into a lipid bilayer, which creates nanopores with typical diameters ranging from ~1-4 nm^68^, although larger pores with diameters up to ~40 nm, e.g. the nuclear pore complex^69^, are also found in nature. Figure 3a shows a schematic of a standard setup for measuring the ionic current through such a protein pore. Briefly, a thick (tens of micrometers) insulating film of amorphous polytetrafluoro-ethylene (PTFE, or Teflon) separates two liquid compartments and contains a ~50-100 µm sized hole where the lipid bilayer is assembled^3,70^. Teflon is the preferred support material due to the relatively low high-frequency noise, and ease of fabrication^71^. Insertion of a protein pore (Fig.3b) short-circuits the insulating bilayer membrane and an ionic current between the two reservoirs can be measured by a pair of Ag/AgCl electrodes. The current signal is amplified by a transimpedance amplifier (e.g. Axopatch 200B) and digitized by an analog-to-digital converter (ADC, e.g. Axon Digidata, same supplier). To shield from external radiative electric noise, the flow-cell and the amplifier headstage are enclosed in a metallic Faraday cage^3^. For biological nanopores, ionic conductances are typically on the order of 0.1-2 nS.

**Figure 3:**
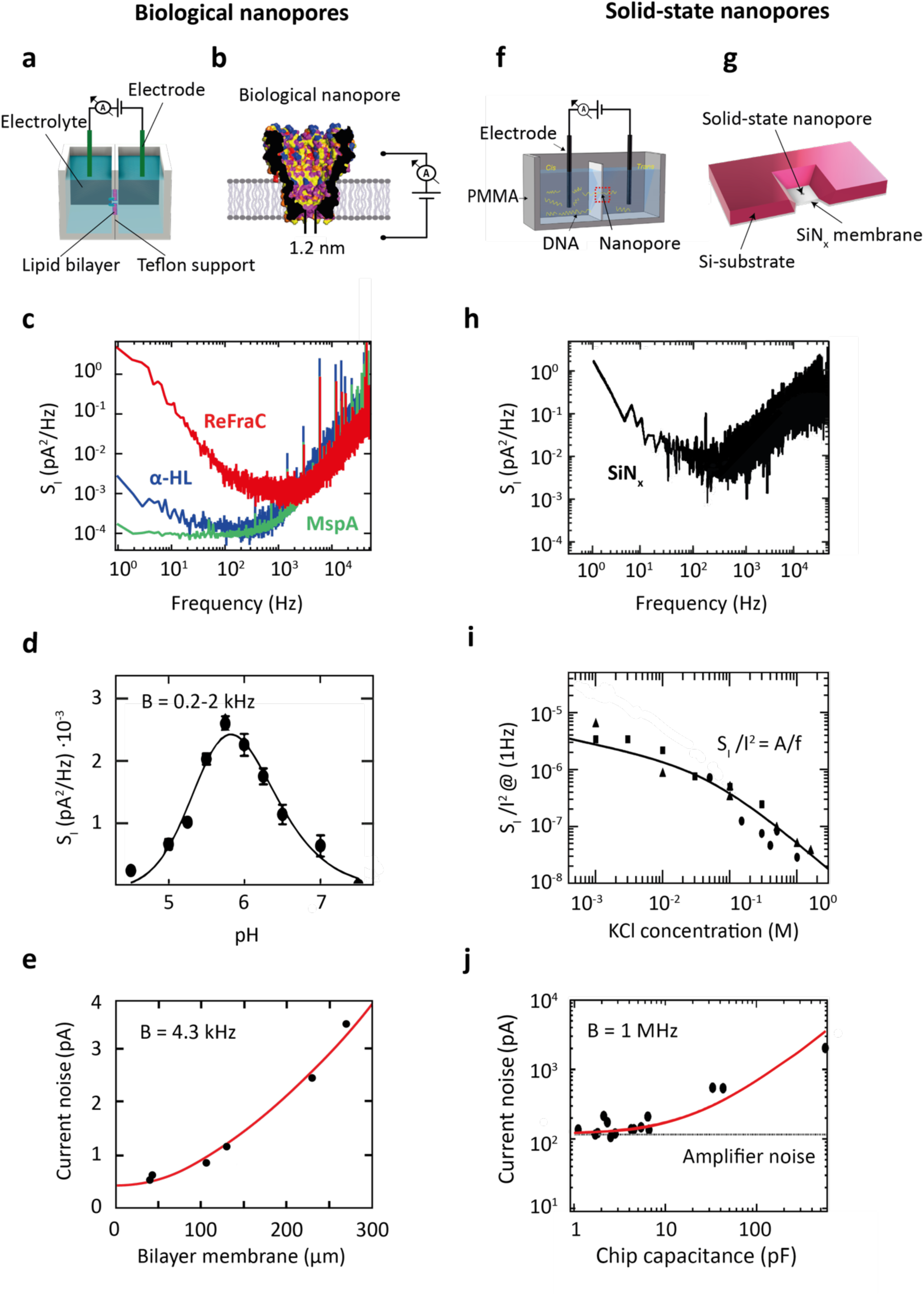
Noise in biological and solid-state nanopores. (a) Standard setup used for measuring the ionic current through a biological nanopore embedded within a lipid membrane. (b) Sketch of a biological MspA nanopore (adapted from Derrington et al., 2010^14^). (c) Typical current PSD for three biological nanopores, ReFraC (D10R/K159E mutant of FraC) ^39^(red), *α*-HL (blue), and the D90N/D91N/D93N/D118R/E139K/D134R mutant of MspA (green), measured in the same setup at 50 mV applied voltage, 1 M KCl salt, pH 7. (d) Low-frequency protonation noise of *α*-HL as a function of pH (adapted from Kasianowicz et al., 1995^61^). (e) Current noise I_rms_ measured at a 4.3 kHz bandwidth of a lipid bilayer setup (where no pore was inserted) vs the size of the bilayer membrane (adapted from Mayer et al., 2003^71^). (f) Schematic of a typical flow cell for measuring the ionic current through a solid-state nanopore (adapted from Feng et al., 2015^4^). (g) Sketch of a solid-state nanopore fabricated onto a Si-supported SiN_x_ membrane. (h) Current PSD for a 15 nm SiN_x_ solid-state nanopore. Data were measured at 100 mV applied voltage for 1 M KCl salt. (i) Relative low-frequency noise S_I_/I^2^ at 1 Hz versus salt concentration. Solid line shows a fit to the data using Hooge’s relation, cf. Eq.(1). (h) and (i) were adapted from Smeets et al., 2008^55^. (j) Current noise I_rms_ measured at a 1 MHz bandwidth vs capacitance of the nanopore chip (adapted from Balan et al., 2014^16^).

Characteristic examples of the current PSD for 3 biological nanopores (α-HL^37^, MspA, and ReFraC^39^) are shown in Figure 3c, as measured at 1 M KCl, pH 7.5, under 50 mV applied bias. Noticeably, both α-HL and MspA exhibit a noise plateau at low frequencies (< 1 kHz) which is due to protonation noise, cf. Eq. (2) for *f* ≪ 1/*τ*. The associated PSD is ~10^−4^ to 10^−3^ pA^2^/Hz, which is higher than the corresponding white noise of ~10^−5^ pA^2^/Hz, set by the sum of thermal and shot noise, Eq. (3) and (4). In the context of single-molecule sensing, protonation noise in biological nanopores was first investigated by Bezrukov and Kasianowicz in the mid 1990s^60,61^. Spectral analysis of the current noise of alpha-hemolysin pores revealed the presence of a Lorentzian-shaped component at low-frequencies (0.2-2 kHz). Given the strong dependence on pH (Fig. 3d), this noise source was associated to the reversible protonation of ionizable residues occurring in the alpha-hemolysin constriction. This notion was further established in a later work by Nestorovich *et al*^59^, where the bacterial porin, OmpF, was shown to produce a similar pH-dependence of the protonation noise.

ReFraC instead shows a pronounced 1/f noise with a PSD of ~10^−1^ pA^2^/Hz at 1 Hz, which is almost three times more than for α-HL and MspA. 1/f noise in biological nanopores was first studied by Benz and coworkers^53,72^, and described using Hooge’s model, Eq.(1). The low-frequency fluctuations observed in a family of bacterial porins were associated with a number of possible phenomena, e.g. gating of the pore channel^53^. In later work by Bezrukov and Winterhalter^54^, conformational changes of the protein pore channel, termed ‘channel breathing’^73^, were discussed as the main cause for the observed 1/f noise.

At higher frequencies (> 1 kHz), the noise in biological nanopores is dominated by dielectric noise arising from the loss conductance of the lipid membrane. In fact, since the dielectric loss and dielectric constant of the teflon are relatively low (*D* = (0.8 − 2) × 10^−4^ and *ε*_*r*_ = 1.89 − 1.93, respectively), the major contribution to the dielectric noise is set by the capacitance of the thin lipid bilayer membrane. This can be attenuated by reducing the area of the teflon hole^71,74^ (Fig. 3e).

### Noise in solid-state nanopores

Solid-state nanopores are generally fabricated in a freestanding membrane of a solid-state material such as silicon nitride (SiN_x_)^75^, graphene^19^, hexagonal boron nitride (h-BN)^76^, or molybdenum disulfide (MoS_2_)^41^, with thicknesses ranging from ~0.3-30 nm. In common nanopore chips (Fig.3g), such a membrane is structurally supported by a ~200-500 µm thick substrate material, typically silicon^75^ (Si), glass^16^ (SiO_2_), or Pyrex^77^. Nanopores can be drilled into the membrane in a variety of ways, e.g. by using a transmission electron microscope (TEM)^78,79^, focused ion beam milling (FIB)^80,81^, reactive ion etching (RIE)^82^, laser-etching^83,84^, or by dielectric breakdown^85,86^, resulting in pore diameters from sub-1nm to tens of nanometers. In a standard solid-state-nanopore experiment, the chip is sandwiched between two rubber O-rings that seal two compartments containing the electrolyte solution (Fig.3f). Alternatively, solid-state pores of ~5-50 nm size can be made by mechanical pulling of hollow glass (SiO_2_) pipettes^87,88^, which are immersed in electrolyte during the measurement. Current sensing, amplification, and recording is the same as for biological nanopores.

Figure 3h displays a typical current PSD measured for a 15 nm diameter SiN_x_ solid-state nanopore^55^ in a 20 nm thick membrane. Characteristic of solid-state nanopores is the pronounced 1/f noise that dominates the low-frequency part of the spectrum (<100 Hz). It can originate from a range of physical processes, see Eq. (1) and associated discussion. Smeets *et al* 2006^49^ showed that poor wettability of the pore surface, associated with the formation of nanobubbles, resulted in high 1/f noise in SiN_x_. Tabard-Cossa *et al*^89^ discussed that high 1/f noise in SiN_x_ pores correlates with surface contamination: inhomogeneities of the pore surface resulted in fluctuations of the number and mobility of charge carriers due to trapping at the pore surface^57,89^, analogous to 1/f noise found in semiconductors^90^. As shown by Smeets *et al*^55,56^, such low-frequency noise in SiN_x_ pores obeys Hooge’s relation, Eq. (1), which describes an inverse proportionality between the 1/f current noise power and the number of charge carriers present within the nanopore volume (Fig.3i)^48^. For nanopores made in 2D materials, the 1/f noise depends strongly on the size of the freestanding area^22,51,52,91^, indicating that mechanical fluctuations of the ultrathin 2D membrane (thickness <1 nm) are the main source. The high-frequency noise in solid-state nanopores is dominated by dielectric (~2-10 kHz) and capacitive noise (>10 kHz)^16,92^, see Fig. 3j. The PSD of these noise sources depends mostly on the capacitance of the chip, cf. Eq. (5) and (6), which in turn is set by the membrane and substrate size, thickness, and dielectric constant. Additionally, parasitic capacitances from the amplifier and the interconnects between nanopore and amplifier contribute to the total capacitance at the amplifier input.

### Comparing the performance of biological and solid-state nanopores

So far, we provided a general overview of the typical noise sources in biological and solid-state nanopores. We now turn to a mutual comparison between these two classes of nanopores. We compare their performances in terms of the SNR – a more relevant figure of merit than the mere magnitude of the current noise. We define the SNR as the ratio between the signal modulation ∆I produced by the translocation of a ssDNA molecule, and the baseline current rms (I_rms_) measured at the operating bandwidth (Fig.4a). Although other definitions of SNR are found in the literature, e.g. as the ratio between open pore current and baseline current noise I_o_/I_rms_^93^ or the capability to discern current levels when sequencing DNA^13,36^, we find this definition the most appropriate for comparison of the relevant translocation signals in a variety of nanopores.

**Figure 4:**
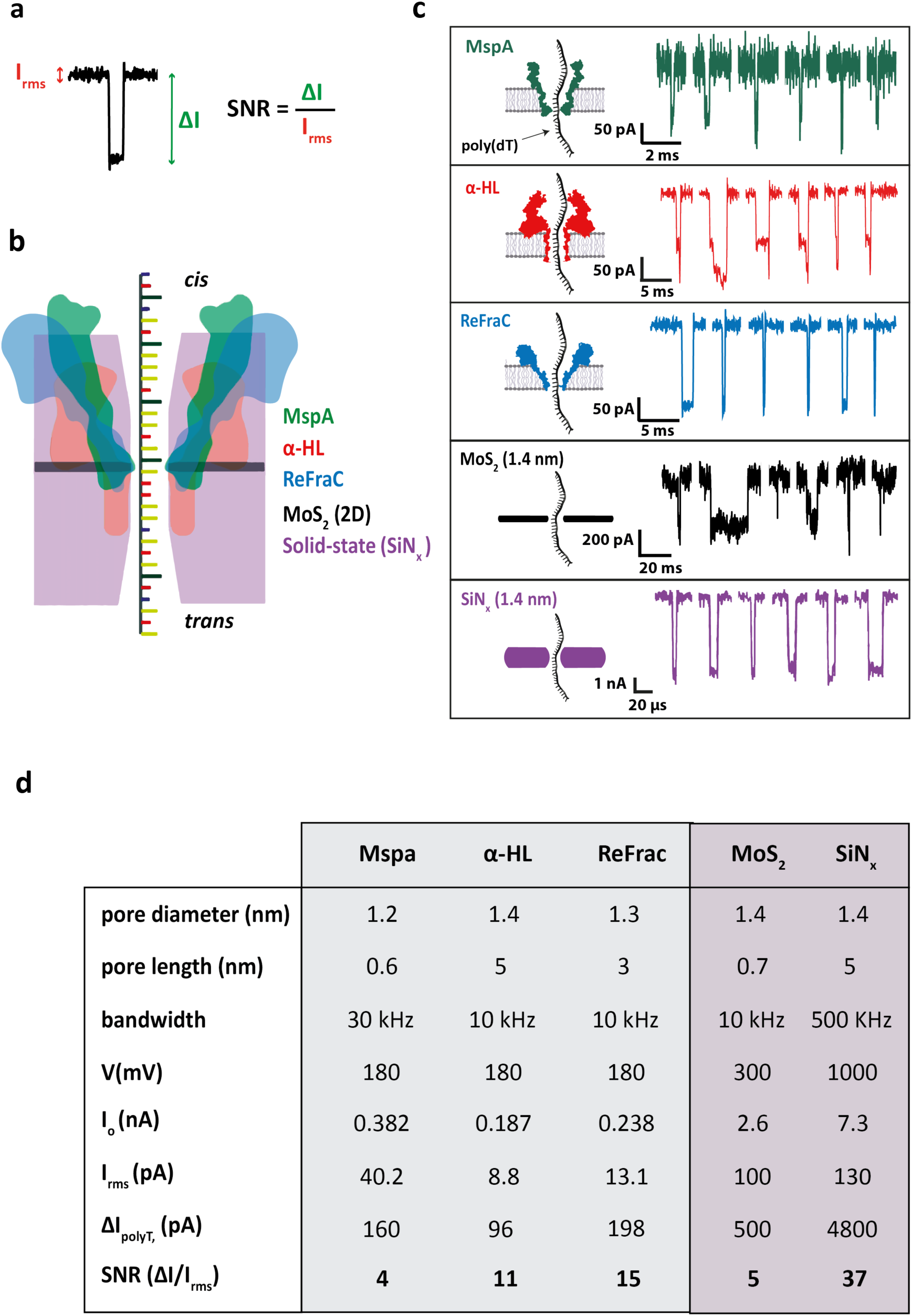
Detection of DNA homopolymer poly(dT) with protein and solid-state nanopores. (a) Example of a translocation event, illustrating the signal-to-noise ratio. (b) Schematic comparing the relative sizes of MspA (green), α-HL (red), ReFraC (blue), MoS_2_ (black), and solid-state SiN_x_ (purple), adapted from Carson & Manunu, 2015^2^. (c) Example of translocation events of poly(dT) molecules through MspA^14^ channel (green), α-HL pore (red), ReFraC pore (blue), 1.4 nm MoS_2_ pore (black), and 1.4 nm SiN_x_ pore (purple, adapted from Venta et al., 2013^18^) all in 1 M KCl solution at a transmembrane voltage of 180 mV, 180 mV, 180 mV, 300mV, and 1V, and at a bandwidth of 30 kHz, 10 kHz, 10 kHz, 10 kHz, and 500 kHz, respectively. Experiments for biological pores were done using an Axopatch 200B amplifier, a teflon-supported lipid membrane (~50-100 µm wide; DPhPC lipids), 10-30 kHz bandwidth, 1 M KCl, pH 7.5, and a forward bias voltage of 180 mV, as in Ref.94. The solid-state SiN_x_ pore was built on a glass chip and measured with the VC100 high-bandwidth, low-noise voltage-clamp amplifier (Chimera Instruments, New York, NY, USA) which allowed for low-noise measurements at high bandwidth. Notably, the positively charged constriction of ReFraC causes the negatively charged poly(dT)_50_ to translocate with much slower (491 ± 114 µs) translocation times compared to MspA (17.7 ± 1.1 µs), which permitted to filter out more high-frequency noise. (d) Comparison of various figures of merit for different nanopore systems under typical experimental conditions. *I*_*O*_ indicates the open pore ionic current at the applied bias V.

Given that the experimental conditions reported in the literature differ considerably, we carried out a dedicated comparative study by complementing reported data with new data that were, to the extent possible, obtained under the same experimental conditions. Bandwidth and applied voltage were chosen such as to fully resolve the current blockade ∆I generated by the poly(dT) substrate (rather than being limited by a too narrow bandwidth). We selected 5 popular nanopore systems, MspA, a-HL, ReFraC, MoS_2_, and SiN_x_, that are commonly used and that were shown to possess good spatiotemporal resolution, allowing for accurate discrimination of short homopolymers^13,14,27,29,39^. All pores considered had a similar diameter of ~1.3 nm. Figure 4b illustrates the relative sizes of the different pores.

Nanopore experiments probing the translocation of poly(dT)_50_ were carried out using three biological pores, MspA, α-HL, and ReFraC. We compared these data to experimental results on two types of solid-state nanopores, SiN_x_^29^ and MoS_2_^27^ that were measured at the same electrolyte conditions. Translocation data of poly(dT)_80_ through a 1.4 nm MoS_2_ pore were kindly shared by the Radenovic lab^41^, whereas poly(dT)_30_ data for a 1.4 nm SiN_x_ pore with ~5 nm length were taken from the literature^29^. Figure 4c shows examples of single-molecule poly(dT) translocations for the 5 pores. A range of SNR values are observed, with, at face value, a better performance for SiN_x_ and ReFraC than for MoS_2_, α-HL, and MspA.

Figure 4d quantitatively compares the data for the different nanopore systems. For the biological nanopores, ReFraC gives the best SNR = 15, while MspA resulted in a much lower SNR of 4. This is mainly due to the faster translocations of poly(dT) through MspA, which required a higher bandwidth (30 kHz), and hence larger noise, in order to resolve the translocation events. Amongst the solid-state nanopores, SiN_x_ showed the best SNR: an impressive value of 37, which was higher than the SNR = 5 obtained for MoS_2_, as well as higher than the values for all biological nanopores. The greater SNR for SiN_x_ results from the very high voltage applied (1000 mV vs 300 mV for MoS_2_), producing a particularly large current signal ∆I. The applied voltage for MoS_2_ pores was limited by the degradation of the 2D membrane and pore growth under high bias voltages, which typically limited the applied bias to < 400 mV. In biological nanopores the range of bias voltages is limited by the membrane stability, affected by electroporation and rupture around 200-300 mV^95,96^. Note furthermore that the SiN_x_ nanopore system was operated at a much higher bandwidth (500 kHz vs 10 kHz for MoS_2_), the regime where dielectric and capacitive noise dominate. This is advantageous for high-voltage sensing, since these noise sources do not scale with voltage, cf. Eq.(5) and Eq.(6). As a result, the high bias voltage improves the signal (∆I) while it does not affect the noise. Lastly, we note that, while MoS_2_ has a lower SNR than SiN_x_, it features a better spatial resolution along the molecule, given its 0.7 nm pore length, as compared to the ~5 nm of SiN_x_.

Finally, it is important to point out that the above comparison was carried out for free translocation of DNA through nanopores. A controlled slowdown of the translocation speed can change these numbers dramatically. Indeed, despite the fact that Figure 4d shows that the best SNR was obtained for the solid-state SiN_x_ nanopores, with values exceeding those of biological nanopores, todays commercialized nanopore-based DNA sequencers employ protein pores to read off DNA bases with (sub)-nucleotide resolution over very long reads^15,97^. Using a helicase to slow down ssDNA molecules through MspA, allowed Laszlo *et al*^36^ to use a very low LP filter frequency of ~200 Hz, and fully resolve the step-wise DNA translocation at half-base resolution. By comparing the noise at a 200 Hz bandwidth with the signal obtained for free poly(dT) translocations in our experiments, we find an exquisite SNR of ~650 for MspA – two orders of magnitude higher than the SNR = 4 noted above. Applying the same reasoning to α-HL and ReFraC increases their SNR to ~270 and ~220, respectively, i.e., somewhat lower values, consistent with their higher low-frequency noise compared MspA (Fig.3c). Thus, in the context of DNA sequencing, the real game-changer lies in the enzymatic control over the translocation speed by use of an additional motor protein^13–15,36,98^. For solid-state nanopores, time control has so far remained a challenge, and accordingly, DNA sequencing has not yet been realized with such nanopores.

### Approaches to overcome noise limitations

Figure 5 shows important approaches to lower the ionic current noise in nanopores. We first describe efforts to reduce the low-frequency noise. As protonation noise is the main source of low-frequency noise in biological nanopores, it is advantageous to choose a pH value that is far away from the pK_a_ of the ionizable amino acids to attenuate the noise. Another way to reduce it, is to remove charged amino acids near the constriction site, which is expected to yield lower noise levels. Furthermore, increasing the conformational stiffness of biological pores can help to reduce conductance fluctuations associated with channel breathing.

**Figure 5:**
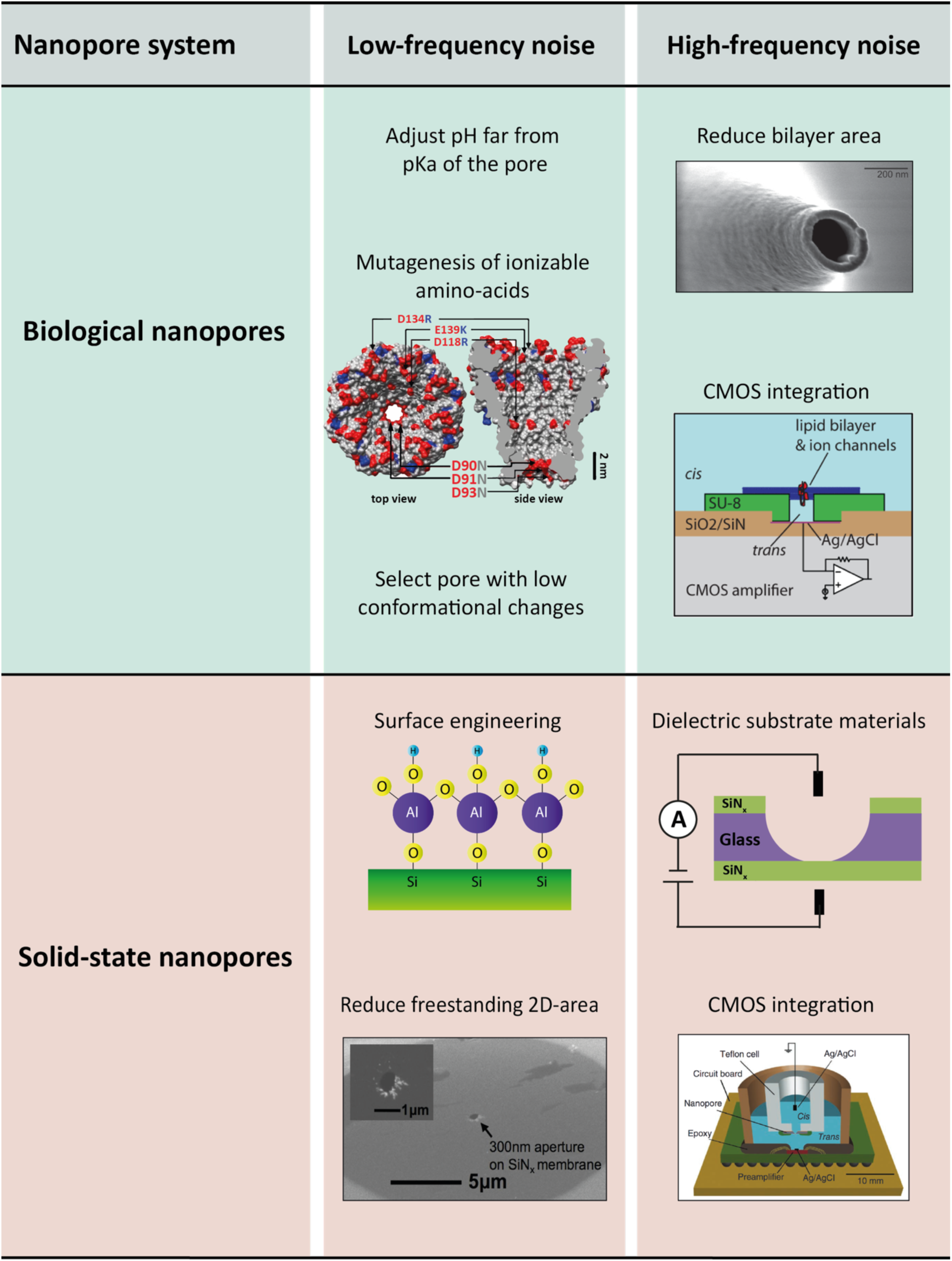
Approaches to reduce the noise in nanopore systems. For biological nanopores, low-frequency protonation noise can be minimized by adjusting the pH far from pKa of the amino acids in the pore constriction, as reported in Ref.60, or by mutating the ionizable amino acids (Arg, Lys, Asp, Glu) to neutral ones (e.g., Asn), as was done for MspA (adapted from Ref.94). Low-frequency 1/f noise instead can only be avoided by selecting a pore that is mechanically stable under an applied bias, e.g. MspA or α-HL. High-frequency noise can be minimized by reducing the size of the freestanding lipid membrane by, e.g., employing a nanocapillary as a support (adapted from Ref.110), and by reducing the capacitance of the interconnects by smart CMOS integration (adapted from Ref.93). For solid-state nanopores, low-frequency 1/f noise can be reduced by coating the surface with a hydrophilic, homogeneous material, e.g. Al_2_O_3_, as reported in Ref.102. For 2D-materials, 1/f noise can be suppressed by lowering the area of the freestanding 2D membrane (adapted from Ref.17). High-frequency noise can be minimized by employing dielectric chip substrate materials, e.g. glass (adapted from Ref.17), or by tight integration of the amplifier and nanopore chip (adapted from Ref.28).

For solid-state nanopores, the low-frequency 1/f noise can be efficiently suppressed by surface functionalization of the SiN_x_ nanopore with a hydrophilic surface layer, such as Al_2_O_3_ or SiO_2_^99–102^. In principle, any surface treatment that reduces the amount of contaminants and improves hydrophilicity of the pore surface will lower the 1/f noise. Indeed, Tabard-Cossa *et al*^89^ showed that piranha treatment (30% H_2_O_2_/H_2_SO_4_, 1:3) substantially reduced the 1/f noise by up to three orders of magnitude. Beamish *et al*^103^ demonstrated that cyclic application of high electric fields to the nanopore also suppressed this noise source. Similar to protein pores, work from Wen *et al*^50^ showed that the 1/f noise could be minimized by choosing a pH that is far from the isoelectric point of the nanopore material (~5 for Si_3_N_4_ ^104,105^). Nanopores built with 2D materials suffer from pronounced 1/f noise that was found to correlate with the area and thickness of the freestanding 2D-membrane^47,48^. A decrease of the freestanding area was shown to reduce the 1/f noise, while employing multi-layer membranes was also helpful for obtaining less noise, though that approach is less desirable due to a loss of spatial resolution^23,51,52^.

The noise at higher frequencies, constituted by dielectric and capacitive noise, has a well-characterized physical origin, namely the thermal voltage noise in conjunction with the loss conductance of the membrane and substrate materials as well as the amplifier input capacitance. Suppression of dielectric noise is generally achieved by minimizing the capacitance *C*_*chip*_ and dielectric loss *D* of the chip, cf. Eq.(5). To effectively decrease capacitive noise, the total input capacitance *C*_*tot*_ needs to be reduced, see Eq.(6) and related discussion. In biological nanopores, the high-frequency noise can be reduced by decreasing the area of the lipid bilayer. Mayer *et* al^71^ fabricated teflon holes of only ~25 µm in diameter with soft lithography using SU-8 resist as master mold, providing a *C*_*chip*_ of 10-28 pF. By using a U-shaped teflon patch tube as the support, Akeson and coworkers^74,94^ built horizontal bilayers < 20 µm in diameter. Lipid bilayers with a comparable size were also created with the droplet-interface-bilayer (DIB) technique^106^. Kitta *et al*^107^ reported on the fabrication of yet smaller bilayers, with sizes down to 2-3 µm in diameter, by using a heated tungsten tip to create a microhole across the teflon film.

Similarly sized 1-3 µm bilayers can be obtained by inserting protein pores into GUVs (Giant Unilamellar Vescicles) and using patch-clamp pipets to measure the conductance of the pores^108,109^. More recently, Gornall *et al*^110^ showed that borosilicate glass nanopipets with diameters as low as 230 nm could be fabricated and used for current recordings on an OmpF protein channel. Hartel *et al*^111^ achieved high-bandwidth (>500 kHz) recordings with biological pores with CMOS-suspended (Complementary Metal-Oxide-Semiconductor) membranes that were built directly over a ~30 µm well on top of a CMOS-amplifier chip. This offered a reduction of the total input capacitance *C*_*tot*_ to < 4 pF and provided a bandwidth as high as 1 MHz and a SNR > 8 at 500 kHz, for detecting the gating of a RyR1 pore (type 1 ryanodine receptor)^111^. Combined with extended β distribution data analysis^112^ (which exploits the characteristics of the excess current noise to reconstruct the true current signal), it was possible to achieve a time resolution of 35 ns^111^.

For reducing the high-frequency noise in SiN_x_ solid-state nanopores, an established method, first reported by Tabard-Cossa *et al*^89^, is to lower *C*_*chip*_ by coating the area of the chip around the pore with a dielectric, e.g. PDMS, thereby providing additional thickness to the chip membrane surrounding the pore and thus a low series capacitance. Similarly, a substantial reduction of *C*_*chip*_ was achieved by employing a dielectric, e.g. amorphous glass^16,17^ or Pyrex^23^ as substrate material instead of the commonly used crystalline silicon which is intrinsically conductive. In work by Balan *et al*^17^, glass chips were shown to reduce *C*_*chip*_ to < 1 pF, compared to > 300 pF for standard silicon chips^55^. Similarly to biological nanopores, the highest working bandwidths were so far achieved by integrating a low *C*_*chip*_ nanopore device with an on-chip CMOS-amplifier^28,30^, which lowered the total input capacitance to *C*_*tot*_ ≈ 4 pF. In this way, ssDNA molecules were recorded using ultrathin (<4 nm) sub-2 nm pores yielding a SNR>10 at 5 MHz^30^. In 2D nanopores, the high-frequency noise can be addressed in similar ways to SiN_x_ pores. The use of glass as substrate material, combined with a small ~300 nm freestanding 2D-membrane of graphene or MoS_2_, resulted in a *C*_*chip*_ < 2 pF^17^.

### Conclusions

In this paper, we illustrated the main sources of noise affecting various nanopore systems, with a particular emphasis on comparing biological and solid-state nanopores, and we discussed practical approaches to lower the noise. We compared the SNR of poly(dT) translocations through a representative set of biological and solid-state pores, and found that silicon nitride nanopores gave the highest SNR. This can be attributed to the higher currents (i.e. larger signals) that solid-state systems offer, and to the relatively low high-frequency noise. Despite these good noise characteristics, prominent applications such as DNA or protein sequencing have so far remained out of reach for solid-state nanopores, because the fast translocation speed provides only a short observation time per single molecule. There are two ways to improve this: one can either shift the sampling rate into even higher frequencies (≫ MHz), or alternatively slow down the translocation of the molecule. The latter strategy has led to the successful commercialization of DNA sequencers based on protein nanopores that are coupled with an enzymatic stepping motor. In our comparison, we found that the SNR of MspA increased >160-fold by such speed control, mainly due to the decoupling of the signal from the high-frequency noise. Additionally, the motor protein provides a ratcheting mechanism that translocates the substrate with a constant discrete step size. Since the sensing region of the pore is typically larger than the individual monomer size (nucleotide or amino-acid), such a mechanism is indispensable to reproducibly resolve and identify the sequence. Future improvements of the solid-state nanopore system could thus be directed towards either a further increase of the temporal resolution, e.g. by reducing even more the overall parasitic capacitances, or by creating an efficient slowdown mechanism, similar to biological nanopores. In general, the understanding of noise sources, associated timescales, and techniques to lower the noise at both low and high frequencies are greatly beneficial to maximize the sensitivity of nanopore detection and thereby extend the range of its applications.

## Acknowledgements

We would like to thank Aleksandra Radenovic, Michael Graf, and Thakur Mukeshchand (EPFL, Switzerland) for sharing and discussing current traces measured on MoS_2_ nanopores; Marija Drndic and Siddharth Shekar (University of Pennsylvania, USA) for sharing current measurement performed on SiN_x_ nanopores; Hagan Bayley and Nicholas Bell (Oxford University, UK) for sharing current traces measured through alpha-hemolysin pores; and Giovanni Maglia and Gang Huang (University of Groningen, the Netherlands) for FraC mutants. Alpha-hemolysin was a kind gift of Jingyue Ju and Sergey Kalachikov (Columbia University, USA); MspA was a kind gift of Jens Gundlach and Andrew H. Laszlo (University of Washington, USA). We thank Meng-yue Wu for technical assistance on TEM, and Wayne Yang, Stephanie Heerema, Laura Restrepo Perez, Sergii Pud, Daniel Verschueren (TU Delft, the Netherlands) for fruitful discussions. This work was supported by ERC Advanced Grant SynDiv (no. 669598) and the NanoFront and BaSyC programs. SS acknowledges the Postdoc.Mobility fellowship no. P400PB_180889 by the Swiss National Science Foundation.

## References

1. Dekker, C. Solid-state nanopores. Nat. Nanotechnol. 2, 209–216 (2007).

2. Carson, S. & Wanunu, M. Challenges in DNA motion control and sequence readout using nanopore devices. Nanotechnology 26, 74004 (2015).

3. Maglia, G., Heron, A. J., Stoddart, D., Japrung, D. & Bayley, H. Analysis of Single Nucleic Acid Molecules with Protein Nanopores. Methods in Enzymology 475, (Elsevier Inc., 2010).

4. Feng, Y., Zhang, Y., Ying, C., Wang, D. & Du, C. Nanopore-based fourth-generation DNA sequencing technology. Genomics, Proteomics Bioinforma. 13, 4–16 (2015).

5. Lin, Y., Ying, Y. L. & Long, Y. T. Nanopore confinement for electrochemical sensing at the single-molecule level. Curr. Opin. Electrochem. 7, 172–178 (2018).

6. Atas, E., Singer, A. & Meller, A. DNA sequencing and bar-coding using solid-state nanopores. Electrophoresis 33, 3437–3447 (2012).

7. Yang, W. et al. Detection of CRISPR-dCas9 on DNA with Solid-State Nanopores. Nano Lett. 18, 6469–6474 (2018).

8. Squires, A. H., Hersey, J. S., Grinstaff, M. W. & Meller, A. A nanopore-nanofiber mesh biosensor to control DNA translocation. J. Am. Chem. Soc. 135, 16304–16307 (2013).

9. Miles, B. N. et al. Single molecule sensing with solid-state nanopores: novel materials, methods, and applications. Chem. Soc. Rev. 42, (2012).

10. Wu, D., Bi, S., Zhang, L. & Yang, J. Single-molecule study of proteins by biological nanopore sensors. Sensors (Switzerland) 14, 18211–18222 (2014).

11. Storm, A. J. et al. Fast DNA translocation through a solid-state nanopore. Nano Lett. 5, 1193–1197 (2005).

12. Plesa, C. et al. Fast translocation of proteins through solid state nanopores. Nano Lett. 13, 658–663 (2013).

13. Manrao, E. a et al. Reading DNA at single-nucleotide resolution with a mutant MspA nanopore and phi29 DNA polymerase. Nat. Biotechnol. 30, 349–353 (2012).

14. Manrao, E. et al. Nanopore DNA sequencing with MspA. Proc. Natl. Acad. Sci. 107, 16060–16065 (2010).

15. Carter, J.-M. & Hussain, S. Robust long-read native DNA sequencing using the ONT CsgG Nanopore system. Wellcome Open Res. 2, 1–11 (2018).

16. Balan, A. et al. Improving signal-to-noise performance for DNA translocation in solid-state nanopores at MHz bandwidths. Nano Lett. 14, 7215–7220 (2014).

17. Balan, A., Chien, C. C., Engelke, R. & Drndic, M. Suspended Solid-state Membranes on Glass Chips with Sub 1-pF Capacitance for Biomolecule Sensing Applications. Sci. Rep. 5, 1–8 (2015).

18. Venta, K. et al. Differentiation of short, single-stranded DNA homopolymers in solid-state nanopores. ACS Nano 7, 4629–4636 (2013).

19. Merchant, C. a. et al. DNA translocation through graphene nanopores. Nano Lett. 10, 2915–2921 (2010).

20. Schneider, G. F. et al. Tailoring the hydrophobicity of graphene for its use as nanopores for DNA translocation. Nat. Commun. 4, 1–7 (2013).

21. Schneider, G. F. et al. DNA translocation through graphene nanopores. Nano Lett. 10, 3163–3167 (2010).

22. Zhou, Z. et al. DNA Translocation through hydrophilic nanopore in hexagonal boron nitride. Sci. Rep. 3, 1–5 (2013).

23. Park, K. B. et al. Noise and sensitivity characteristics of solid-state nanopores with a boron nitride 2-D membrane on a pyrex substrate. Nanoscale 8, 5755–5763 (2016).

24. Liu, K. et al. Geometrical Effect in 2D Nanopores. Nano Lett. 17, 4223–4230 (2017).

25. Riaz, N., Wolden, S. L., Gelblum, D. Y. & Eric, J. HHS Public Access. 118, 6072–6078 (2016).

26. Liu, K., Feng, J., Kis, A. & Radenovic, A. Atomically thin molybdenum disulfide nanopores with high sensitivity for dna translocation. ACS Nano 8, 2504–2511 (2014).

27. Feng, J. et al. Identification of single nucleotides in MoS 2 nanopores. Nat. Nanotechnol. 10, 1070–1076 (2015).

28. Rosenstein, J. K., Wanunu, M., Merchant, C. A., Drndic, M. & Shepard, K. L. Integrated nanopore sensing platform with sub-microsecond temporal resolution. Nat. Methods 9, 487–492 (2012).

29. Venta, K. et al. Differentiation of short, single-stranded DNA homopolymers in solid-state nanopores. ACS Nano 7, 4629–4636 (2013).

30. Shekar, S. et al. Measurement of DNA translocation dynamics in a solid-state nanopore at 100 ns temporal resolution. Nano Lett. 16, 4483–4489 (2016).

31. Thiel, G. et al. High bandwidth approaches in nanopore and ion channel recordings – A tutorial review. Anal. Chim. Acta 1061, (2019).

32. Keyser, U. F. Controlling molecular transport through nanopores. J. R. Soc. Interfac. 8, 1369–1378 (2011).

33. Wanunu, M. & Meller, A. Chemically modified solid-state nanopores. Nano Lett. 7, 1580–1585 (2007).

34. Pud, S. et al. Mechanical Trapping of DNA in a Double-Nanopore System. Nano Lett. 16, 8021–8028 (2016).

35. Gilboa, T. & Meller, A. Optical sensing and analyte manipulation in solid-state nanopores. Analyst 140, 4733–4747 (2015).

36. Laszlo, A. H., Derrington, I. M. & Gundlach, J. H. MspA nanopore as a single-molecule tool: From sequencing to SPRNT. Methods 105, 75–89 (2016).

37. Song, L. et al. Structure of staphylococcal α-hemolysin, a heptameric transmembrane pore. Science (80-.). 274, 1859–1866 (1996).

38. Menestrina, G. Ionic Channels Formed by Staphylococcus aureus Alpha-Toxin: Voltage-Dependent Inhibition by Divalent and Trivalent Cations. J. Membr. Biol. 90, 177–190 (1986).

39. Wloka, C., Mutter, N. L., Soskine, M. & Maglia, G. Alpha-Helical Fragaceatoxin C Nanopore Engineered for Double-Stranded and Single-Stranded Nucleic Acid Analysis. Angew. Chemie - Int. Ed. 55, 12494–12498 (2016).

40. Huang, G., Willems, K., Soskine, M., Wloka, C. & Maglia, G. Electro-osmotic capture and ionic discrimination of peptide and protein biomarkers with FraC nanopores. Nat. Commun. 8, 1–13 (2017).

41. Graf, M. et al. Fabrication and practical applications of molybdenum disulfide nanopores. Nat. Protoc. 14, 1130–1168 (2019).

42. Sakmann, B. & Neher, E. Single-channel recording. (2009). doi:10.1007/978-1-4419-1229-9

43. Tabard-Cossa, V. Instrumentation for Low-Noise High-Bandwidth Nanopore Recording. i. Engineered Nanopores for Bioanalytical Applications: A Volume in Micro and Nano Technologies 59–93 (2013). doi:10.1016/B978-1-4377-3473-7.00003-0

44. Milotti, E. 1/F Noise: a Pedagogical Review. arXiv (2002).

45. Dutta, P. & Horn, P. M. Low-frequency fluctuations in solids: 1/f noise. Rev. Mod. Phys. 53, 497–516 (1981).

46. Zhang, D., Solomon, P., Zhang, S. L. & Zhang, Z. An impedance model for the low-frequency noise originating from the dynamic hydrogen ion reactivity at the solid/liquid interface. Sensors Actuators, B Chem. 254, 363–369 (2018).

47. Jindal, R. P. & Van Der Ziel, A. Model for mobility fluctuation 1/f noise. Appl. Phys. Lett. 38, 290–291 (1981).

48. Hooge, F. N. 1/F Noise. Phys. B+C 83, 14–23 (1976).

49. Smeets, R. M. M., Keyser, U. F., Wu, M. Y., Dekker, N. H. & Dekker, C. Nanobubbles in solid-state nanopores. Phys. Rev. Lett. 97, 1–4 (2006).

50. Wen, C. et al. Generalized Noise Study of Solid-State Nanopores at Low Frequencies. ACS Sensors 2, 300–307 (2017).

51. Heerema, S. J. et al. 1/F Noise in Graphene Nanopores. Nanotechnology 26, (2015).

52. Zhang, Z.-Y. et al. Noise Analysis of Monolayer Graphene Nanopores. Int. J. Mol. Sci. 19, 2639 (2018).

53. Wohnsland, F. & Benz, R. 1/f-Noise of open bacterial porin channels. J. Membr. Biol. 158, 77–85 (1997).

54. Bezrukov, S. M. & Winterhalter, M. Examining noise sources at the single-molecule level: 1/f noise of an open maltoporin channel. Phys. Rev. Lett. 85, 202–205 (2000).

55. Smeets, R. M. M., Keyser, U. F., Dekker, N. H. & Dekker, C. Noise in solid-state nanopores. Proc. Natl. Acad. Sci. 105, 417–421 (2008).

56. Smeets, R. M. M., Dekker, N. H. & Dekker, C. Low-frequency noise in solid-state nanopores. Nanotechnology 20, (2009).

57. Fragasso, A., Pud, S. & Dekker, C. 1/F Noise in Solid-State Nanopores Is Governed By Access and Surface Regions. Nanotechnology 30, 395202 (2019).

58. Tasserit, C., Koutsioubas, A., Lairez, D., Zalczer, G. & Clochard, M. C. Pink noise of ionic conductance through single artificial nanopores revisited. Phys. Rev. Lett. 105, 1–4 (2010).

59. Nestorovich, E. M., Rostovtseva, T. K. & Bezrukov, S. M. Residue Ionization and Ion Transport through OmpF Channels. Biophys. J. 85, 3718–3729 (2003).

60. Kasianowicz, J. J. & Bezrukov, S. M. Current Noise Reveals Protonation Kinetics and Number of Ionizable Sites in an Open Protein Ion Channel. Phys. Rev. Lett. 70, 2352–2355 (1993).

61. Kasianowicz, J. J. & Bezrukov, S. M. Protonation dynamics of the alpha-toxin ion channel from spectral analysis of pH-dependent current fluctuations. Biophys. J. 69, 94–105 (1995).

62. Machlup, S. Noise in semiconductors: Spectrum of a two-parameter random signal. J. Appl. Phys. 25, 341–343 (1954).

63. Johnson, J. B. Thermal agitation of electricity in conductors. Phys. Rev. 32, (1928).

64. Nyquist, H. Thermal agitation of electric charge in conductors. Phys. Rev. 32, (1928).

65. Blanter, Y. M. & Büttiker, M. Shot noise in mesoscopic conductors. Phys. Rep. 336, 1–166 (2000).

66. Schottky, W. Über spontane Stromschwankungen in verschiedenen Elektrizitätsleitern. Ann. Phys. 362, 541–567 (1918).

67. Sherman-Gold, R. The Axon CNS Guide to Electrophysiology and Biophysics Laboratory Techniques. (2012).

68. Ayub, M. & Bayley, H. Engineered transmembrane pores. Curr. Opin. Chem. Biol. 34, 117–126 (2016).

69. Yu, Z. et al. Integrative structure and functional anatomy of a nuclear pore complex. Nature 555, 475–482 (2018).

70. Robertson, J. W. F. Biological Pores on Lipid Bilayers. Engineered Nanopores for Bioanalytical Applications: A Volume in Micro and Nano Technologies (Elsevier Inc., 2013). doi:10.1016/B978-1-4377-3473-7.00004-2

71. Mayer, M., Kriebel, J. K., Tosteson, M. T. & Whitesides, G. M. Microfabricated Teflon membranes for low-noise recordings of ion channels in planar lipid bilayers. Biophys. J. 85, 2684–2695 (2003).

72. Nekolla, S., Andersen, C. & Benz, R. Noise analysis of ion current through the open and the sugar-induced closed state of the LamB channel of Escherichia coli outer membrane: evaluation of the sugar binding kinetics to the channel interior. Biophys. J. 66, 1388–1397 (1994).

73. Läuger, P. Structural fluctuations and current noise of ionic channels. Biophys. J. 48, 369–373 (1985).

74. Akeson, M., Brandin, E., Branton, D., Deamer, D. W. & Kasianowicz, J. J. Microsecond Time-Scale Discrimination Among Polycytidylic Acid, Polyadenylic Acid, and Polyuridylic Acid as Homopolymers or as Segments Within Single RNA Molecules. Biophys. J. 77, 3227–3233 (2009).

75. Gibb, T. & Ayub, M. Solid-State Nanopore Fabrication. Eng. Nanopores Bioanal. Appl. A Vol. Micro Nano Technol. 121–140 (2013). doi:10.1016/B978-1-4377-3473-7.00005-4

76. Gilbert, S. M. et al. Fabrication of Subnanometer-Precision Nanopores in Hexagonal Boron Nitride. Sci. Rep. 7, 1–7 (2017).

77. Lee, M. H. et al. A low-noise solid-state nanopore platform based on a highly insulating substrate. Sci. Rep. 4, 1–7 (2014).

78. Storm, A. J., Chen, J. H., Ling, X. S., Zandbergen, H. W. & Dekker, C. Fabrication of solid-state nanopores with single-nanometre precision. Nat. Mater. 2, 537–540 (2003).

79. Van Den Hout, M. et al. Controlling nanopore size, shape and stability. Nanotechnology 21, (2010).

80. Lanyon, Y. H. et al. Fabrication of nanopore array electrodes by focused ion beam milling. Anal. Chem. 79, 3048–3055 (2007).

81. Schiedt, B. et al. Direct FIB fabrication and integration of ‘single nanopore devices’ for the manipulation of macromolecules. Microelectron. Eng. 87, 1300–1303 (2010).

82. Verschueren, D. V., Yang, W. & Dekker, C. Lithography-based fabrication of nanopore arrays in freestanding SiN and graphene membranes. 29, 145302 (2018).

83. Gilboa, T., Zvuloni, E., Zrehen, A., Squires, A. H. & Meller, A. Automated, Ultra-Fast Laser-Drilling of Nanometer Scale Pores and Nanopore Arrays in Aqueous Solutions. Adv. Funct. Mater. 1900642, 1–9 (2019).

84. Gilboa, T., Zrehen, A., Girsault, A. & Meller, A. Optically-Monitored Nanopore Fabrication Using a Focused Laser Beam. Sci. Rep. 8, 1–10 (2018).

85. Pud, S. et al. Self-Aligned Plasmonic Nanopores by Optically Controlled Dielectric Breakdown. Nano Lett. 15, 7112–7117 (2015).

86. Kwok, H., Briggs, K. & Tabard-Cossa, V. Nanopore fabrication by controlled dielectric breakdown. PLoS One 9, (2014).

87. Xu, X., Li, C., Zhou, Y. & Jin, Y. Controllable Shrinking of Glass Capillary Nanopores Down to sub-10 nm by Wet-Chemical Silanization for Signal-Enhanced DNA Translocation. ACS Sensors 2, 1452–1457 (2017).

88. Bafna, J. A. & Soni, G. V. Fabrication of Low Noise Borosilicate Glass Nanopores for Single Molecule Sensing. PLoS One 11, e0157399 (2016).

89. Tabard-Cossa, V., Trivedi, D., Wiggin, M., Jetha, N. N. & Marziali, A. Noise analysis and reduction in solid-state nanopores. Nanotechnology 18, (2007).

90. Vandamme, L. K. J. & Rigaud, D. 1/f noise in MOS devices, mobility or number fluctuations. IEEE Trans. Electron Devices 41, 1936–1945 (1994).

91. Garaj, S., Liu, S., Golovchenko, J. A. & Branton, D. Molecule-hugging graphene nanopores. Proc. Natl. Acad. Sci. U. S. A. 110, 12192–12196 (2013).

92. Roelen, Z., Bustamante, J. A., Carlsen, A., Baker-Murray, A. & Tabard-Cossa, V. Instrumentation for low noise nanopore-based ionic current recording under laser illumination. Rev. Sci. Instrum. 89, (2018).

93. Rosenstein, J. K., Ramakrishnan, S., Roseman, J. & Shepard, K. L. Single ion channel recordings with CMOS-anchored lipid membranes. Nano Lett. 13, 2682–2686 (2013).

94. Butler, T. Z., Pavlenok, M., Derrington, I. M., Niederweis, M. & Gundlach, J. H. Single-molecule DNA detection with an engineered MspA protein nanopore. Proc. Natl. Acad. Sci. 105, 20647–20652 (2008).

95. Pavlin, M., Kotnik, T., Miklavčič, D., Kramar, P. & Maček Lebar, A. Electroporation of Planar Lipid Bilayers and Membranes. in Advances in Planar Lipid Bilayers and Liposomes 6, 165–226 (2008).

96. Tarek, M. Membrane electroporation: A molecular dynamics simulation. Biophys. J. 88, 4045–4053 (2005).

97. Jain, M. et al. Nanopore sequencing and assembly of a human genome with ultra-long reads. Nat. Biotechnol. 36, 338–345 (2018).

98. Jain, M., Olsen, H. E., Paten, B. & Akeson, M. The Oxford Nanopore MinION: delivery of nanopore sequencing to the genomics community. Genome Biol. 17, 1–11 (2016).

99. Wang, C. M., Kong, D. L., Chen, Q. & Xue, J. M. Surface engineering of synthetic nanopores by atomic layer deposition and their applications. Front. Mater. Sci. 7, 335–349 (2013).

100. Nilsson, J., Lee, J. R. I., Ratto, T. V. & Létant, S. E. Localized functionalization of single nanopores. Adv. Mater. 18, 427–431 (2006).

101. Danelon, C., Santschi, C., Brugger, J. & Vogel, H. Fabrication and functionalization of nanochannels by electron-beam-induced silicon oxide deposition. Langmuir 22, 10711–10715 (2006).

102. Chen, P. et al. Atomic layer deposition to fine-tune the surface properties and diameters of fabricated nanopores. Nano Lett. 4, 1333–1337 (2004).

103. Beamish, E., Kwok, H., Tabard-Cossa, V. & Godin, M. Precise control of the size and noise of solid-state nanopores using high electric fields. Nanotechnology 23, (2012).

104. Kosmulski, M. The pH-dependent surface charging and the points of zero charge. J. Colloid Interface Sci. 253, 77–87 (2002).

105. Firnkes, M., Pedone, D., Knezevic, J., Döblinger, M. & Rant, U. Electrically facilitated translocations of proteins through silicon nitride nanopores: Conjoint and competitive action of diffusion, electrophoresis, and electroosmosis. Nano Lett. 10, 2162–2167 (2010).

106. Bayley, H. et al. Droplet interface bilayers RID B-8725-2008. Mol. Biosyst. 4, 1191–1208 (2008).

107. Kitta, M., Tanaka, H. & Kawai, T. Rapid fabrication of Teflon micropores for artificial lipid bilayer formation. Biosens. Bioelectron. 25, 931–934 (2009).

108. Criado, M. & Keller, B. U. A membrane fusion strategy for single-channel recordings of membranes usually non-accessible to patch-clamp pipette electrodes. FEBS Lett. 224, 172–176 (1987).

109. Riquelme, G., Lopez, E., Garcia-segura, L. M., Ferragut, J. A. & Gonzalez-ros, J. M. Giant Liposomes: A Model System in Which To Obtain Patch-Clamp Recordings. Society 11215–11222 (1990).

110. Gornall, J. L. et al. Simple reconstitution of protein pores in nano lipid bilayers. Nano Lett. 11, 3334–3340 (2011).

111. Hartel, A. J. W. et al. Single-channel recordings of RyR1 at microsecond resolution in CMOS-suspended membranes. Proc. Natl. Acad. Sci. 115, E1789–E1798 (2018).

112. Schroeder, I. How to resolve microsecond current fluctuations in single ion channels: The power of beta distributions. Channels 9, 262–280 (2015).

